# Reward-tethered place cells support flexible magnitude coding and remapping in the hippocampus

**DOI:** 10.64898/2026.05.15.725501

**Authors:** Nicola Masala, Margaret M. Donahue, Brittney L. Boublil, Gimarie Irizarry Martinez, Habiba Abouelatta, Kylee B. Miller, Marta Sabariego, Laura A. Ewell

## Abstract

Learning to navigate a changing environment requires the ability to detect when a reward no longer matches a previous expectation. While the hippocampus is essential for adapting behavior strategies when reward expectations are violated and is known to integrate goal-related information into spatial maps, the precise dynamics by which these maps update during an unexpected reduction in reward magnitude is not well understood. Using longitudinal calcium imaging of neuronal activity in behaving mice, we found that hippocampal CA1 population activity encodes reward magnitude through elevated event rates at high value locations. Within this population, we discovered a specialized group of reward tethered place cells that bind spatial context to reward magnitude. These reward tethered place cells exhibit spatial fields across the environment while simultaneously exhibiting activity anchored to high-value reward locations. Upon reward reduction, CA1 population activity equalizes and these neurons undergo a selective and rapid remapping that precedes behavioral adjustment to the reward downshift. The broader spatial map remains intact, indicating that this change allows the animal to update the value of a goal while preserving a stable representation of its surroundings. This selective reorganization of hippocampal firing patterns could support adaptive decision making by updating internal models of the world when expectations are violated.

Graphical abstract.
Masala N., Donahue M., etal.

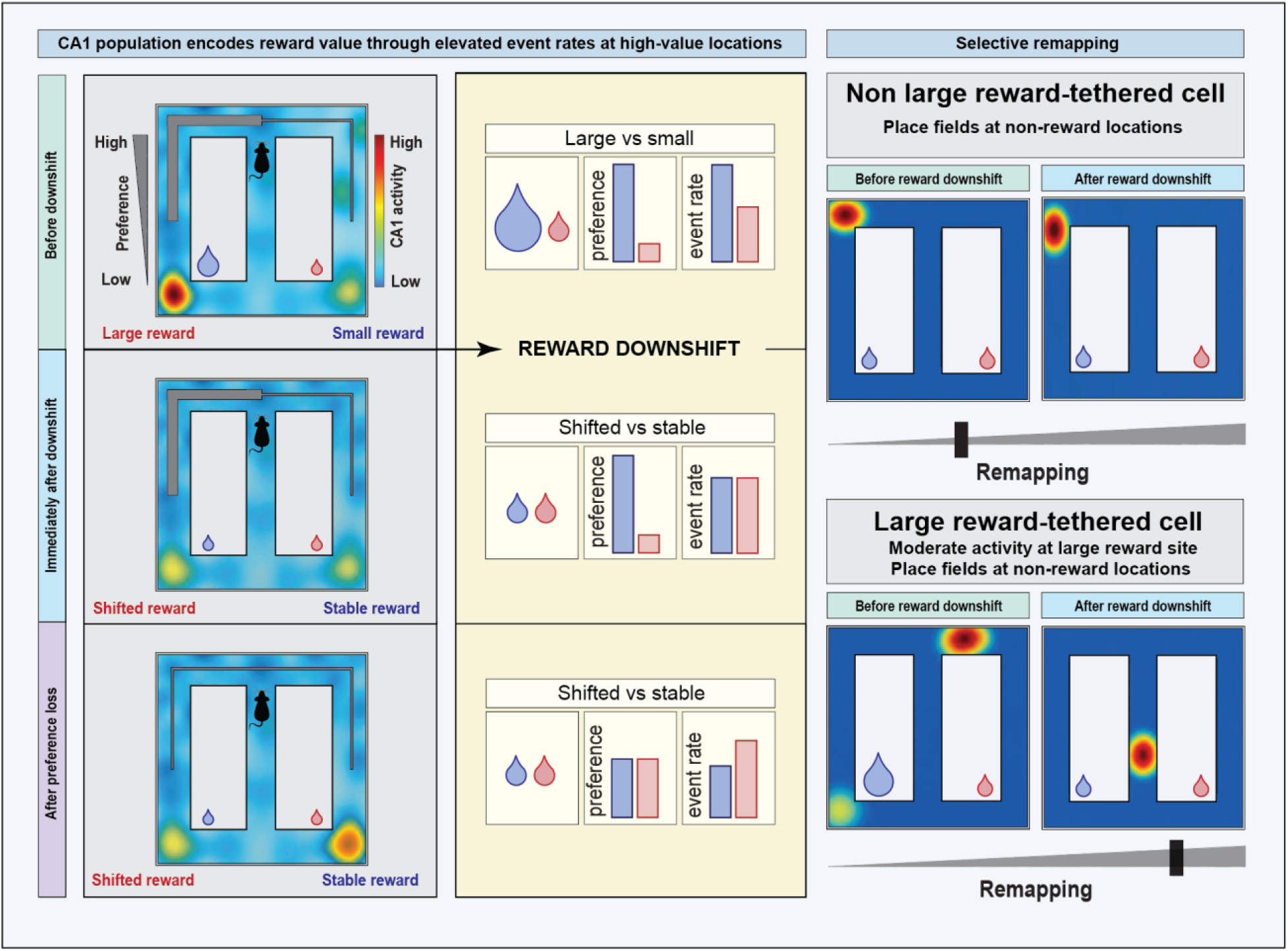

## INTRODUCTION

Survival in a dynamic environment requires the ability to anticipate reward outcomes across distinct locations and to adjust behavior when expectations are not met. The dorsal hippocampus is canonically viewed as the substrate for the cognitive map ^1^, and a growing body of work shows that hippocampal activity is multiplexed, associating spatial environmental features with attention, memory demands, and reward value or magnitude ^2–7^. During learning, hippocampal reward representations undergo a backward shift, transforming from a code of reward consumption to one of reward prediction ^8,7^. This transformation sets the stage for the hippocampus to detect discrepancies between expected and realized outcomes ^9^. Unexpected violations of reward expectancy generate a negative prediction error, which is a signal that triggers a cognitive update of environmental contingencies ^10^. While the behavioral adjustments to changes in reward value have been described for decades ^11–13^, the precise neural dynamics by which the hippocampus updates spatial and non-spatial representations to reflect reward devaluation remain a subject of intense investigation.

Crucially, neural signals related to outcome evaluation, reward magnitude, and prediction error appear to be significantly stronger in the CA1 subregion compared to the upstream area CA3 ^14,15^. Furthermore, the stability of the CA1 place cell code itself appears to be modulated by reward. High-value expectations can stabilize representational drift and ensure that memories of salient goals persist over time, while the loss of reward during extinction can trigger a rapid degradation of spatial maps ^16^. This neural flexibility facilitates the adaptive switch required when expectations no longer match environmental outcomes. Hippocampal representations are highly flexible and change their firing patterns, or “remap”, in response to changes in task demands or reward contingency to ensure efficient navigation toward salient goals ^17,18^.

Despite this progress, a distinction should be made between total reward omission (extinction) and an unexpected reduction in reward magnitude (downshift). While omission involves the total absence of an outcome, a downshift in magnitude presents the system with an approach-avoidance conflict ^19^, triggering a rapid recalibration of goal-value while the animal remains engaged with the environment. This conflict could serve as a cognitive driver, prompting the hippocampal map to resolve the prediction error by remapping to include new predictive cues. In this scenario, the reward is present, but its diminished value violates the magnitude prediction, leading to a process of incentive devaluation. Although we have gained significant insight into how the dorsal hippocampus encodes the presence of reward, a fundamental gap remains in our understanding of how hippocampal representations reorganize specifically following an unexpected downshift in reward magnitude. Building on our previous finding that behavioral adjustment to reward downshift depends on the hippocampus ^20^, we here employed longitudinal calcium imaging to track neuronal activity during this transition. We identified a population-level magnitude code for unequal rewards in CA1, supported by a distinct neuronal class: ‘large reward-tethered place cells’ (LRTs). These cells are characterized by spatial fields distributed throughout the maze and moderate activity at high value reward locations. This population-level magnitude code exhibits rapid plasticity, realigning immediately following reward downshift. Furthermore, the observation that remapping is largely driven by LRTs suggests that these cells could serve as a specialized substrate for resolving prediction errors, integrating spatial context with shifting reward values to update the internal model.

## RESULTS

### Unexpected reward downshift recruits dorsal CA1 and triggers immediate changes in foraging behavior

Our previous work demonstrates that the dorsal hippocampus is necessary for rats to update their foraging behavior in response to an unexpected downshift of reward magnitude ^20^. To determine whether dorsal CA1 is similarly involved in mice, we trained animals on an instrumental reward downshift task in a figure-8 maze. Mice were trained to traverse a central stem and choose between two distinct reward locations that offered unequal magnitudes (Fig. 1 & S1A, B). Over several training sessions, mice developed a consistent preference for the large reward location (Figure 1C & S1C). Once a stable preference was established, we unexpectedly downshifted the large reward to match the small reward magnitude. We found that this unexpected downshift in reward magnitude was associated with a significant increase of cFos-positive cells within dorsal CA1 (Extended Data Fig.1; mean ± SEM; Downshift group (n = 9): 1421 ± 173.4 cells/mm^2^; Unshifted control group (n = 8): 940 ± 109.6 cells/mm^2^, Mann-Whitney, U = 14, p = 0.036).

**Figure 1.**
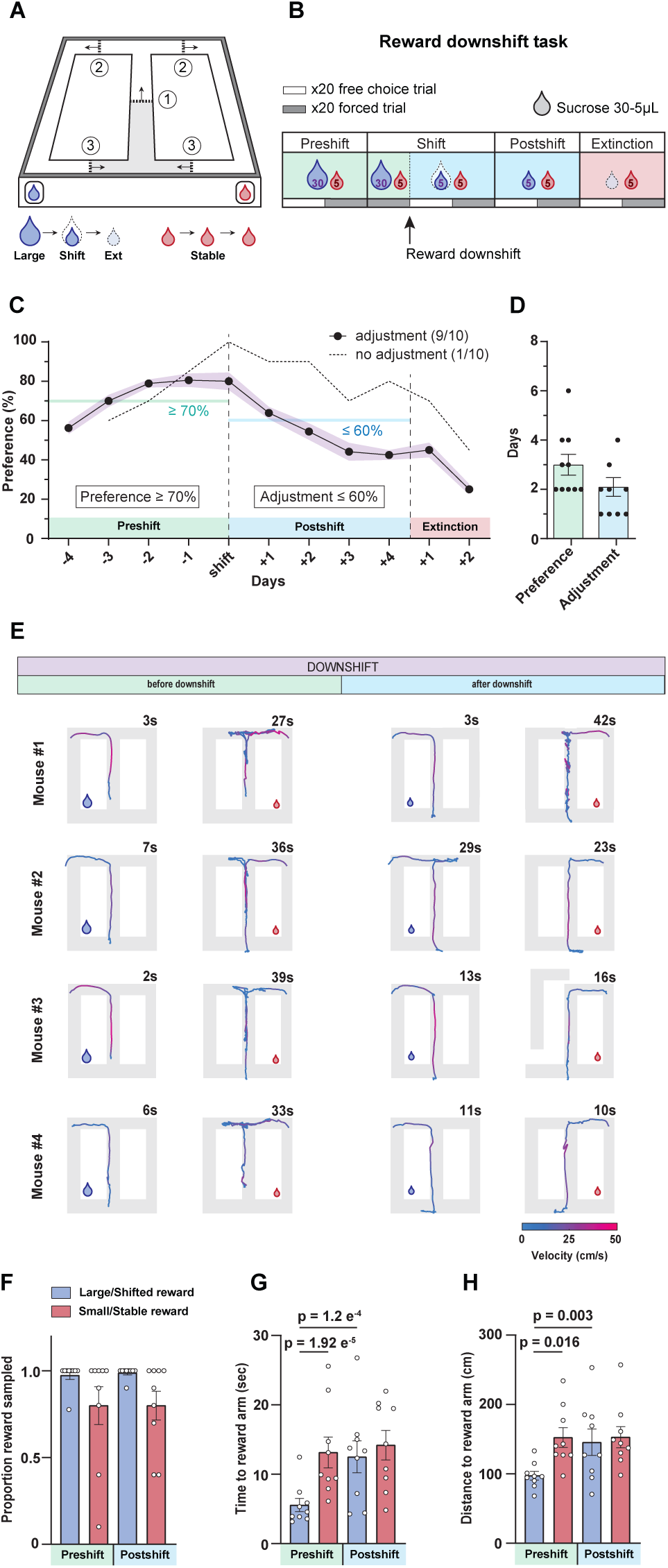
Mice develop a preference for large rewards and exhibit behavioral adjustment after reward downshift. (A) Schematic representation of the automated figure-8 maze for implementing the reward downshift task. The maze is equipped with automatic doors (dotted lines) and reward ports (droplets). Automated doors allowed for free-choice or guided force trials. Mice are held in the delay zone (shaded region) for 15 seconds in between trials. (B) The multi-day task was structured such that mice experienced several reward contingencies: Preshift (unequal rewards of 30 and 5 mL sugar water drops), Shift (large reward is downshifted to match magnitude of small reward), Postshift (equal rewards), and Extinction. Within each of these trial phases there are blocks of free-choice trials and blocks of guided ‘forced’ trials (grey and white key). (C) The behavioral summary (n=9 mice, mean ± sem) for the free-choice blocks show that over several days of the Preshift contingency mice develop a preference for visiting the reward port with the large magnitude reward and subsequently abandon that preference during the Postshift phase. Preference for the large reward was defined as ≥ 70% free-choice trials toward the larger reward port on 2 of 3 consecutive days. Adjustment was defined as ≤ 60% free-choice trials toward the downshifted reward port. One animal did not adjust after downshifting (dotted line) and was excluded from further analysis and calcium imaging experiments. (D) Days to behavioral criterion during preference acquisition and after reward downshift. Light green columns indicate the number of days required for mice to reach the criterion for large-reward preference (≥70%). Light blue columns indicate the number of days required to exhibit loss of preference following large-reward downshift (≤60%). Points represent individual animals. Mice reached preference criterion in 3.0 ± 0.421 days (mean ± SEM) and exhibited behavioral adjustment in 1.8 ± 0.351 days. (E) Example trajectory paths from four representative mice during the downshift day, shown for both reward conditions (large vs. small; downshifted vs. stable) and task phases (preshift and postshift). Time to reward-arm entry is indicated (upper right). Trajectories are color-coded by running velocity, ranging from low (blue, 0 cm s⁻¹) to high (red, 50 cm s⁻¹). (F) Proportion of rewards sampled did not differ between phases (forced trials before vs. after reward downshift). Rewards were more frequently skipped on the small/stable reward side across phases. P-value shown for post-hoc pairwise comparisons following two-way ANOVA (phase x reward). (G) Latency to enter the large-reward arm differed across phases, whereas latency to enter the small-reward arm did not. Mice entered the large-reward arm more rapidly during forced-choice trials before downshift compared to after downshift, with no corresponding phase difference for the small-reward arm. No difference was observed between latencies to enter the downshifted and stable reward arms during post-downshift trials P-values shown for post-hoc pairwise comparisons following two-way ANOVA (phase x reward). (H) Path length to the large-reward arm differed across phases, whereas path length to the small-reward arm did not. Mice traveled shorter paths to the large-reward arm during forced-choice trials before downshift compared to after downshift, with no corresponding phase difference for the small-reward arm. No difference was observed between path lengths to the downshifted and stable reward arms during postshift trials. P-values shown for post-hoc pairwise comparisons following two-way ANOVA (phase x reward).

Given the dorsal CA1 engagement during the reward downshift, we adapted the task to a fully automated version designed for *in vivo* physiology. This version enabled us to monitor CA1 dynamics across all task phases (Fig. 1A, B). To ensure equal sampling of less preferred locations, we added blocks of forced trials in which mice were directed to reward locations via automated doors. The full task comprised three phases: Preshift, Downshift, and Postshift (Fig. 1B, C). In the Preshift phase, reward magnitudes were unequal. Mice remained in this phase until they showed a preference of at least 70 % for the large reward during the free-choice trials. All mice (n = 10; 5 males and 5 females) met this criterion (mean preference: 81 % ± 3.06; Fig. 1C) within 3.0 ± 0.42 days (Fig. 1D). The Downshift phase took place over one day and included three distinct trial blocks: a block of 20 forced trials, a block of 20 free-choice trials, and another block of 20 forced trials. The first block of forced trials matched the Preshift reward contingencies to provide a baseline for the day. During the free choice and second block of forced trials, the magnitude of the large reward was decreased so that both rewards were equal in magnitude (i.e., the large reward was downshifted and the small reward remained stable). Following the Downshift day, the Postshift phase continued until animals exhibited behavioral adjustment, defined as ≤ 60 % preference for the shifted reward during free-choice trials. Nine mice (n = 5 male, 4 female) reached this adjustment within 1.8 ± 0.35 days of the Downshift day (mean preference: 59 % ± 2.47; Fig. 1C). One female mouse maintained an 80 % preference after four days and was excluded from physiological analysis. Following completion of the three task phases, we extinguished the downshifted reward for two additional days, while the stable reward remained available.

Although choice preference remained stable on the day of the downshift (Fig. 1C, Downshift vs. −1), mice exhibited rapid changes in foraging behavior during force trials that were reflected in their trajectory paths (Fig. 1G, H). These effects occurred while overall task engagement remained high and the total proportion of rewards sampled did not change between experimental phases across the two arms of the figure-8 maze (Fig. 1F; n = 9 mice; two way ANOVA, no interaction between phase and reward, F(1,32) = 0.009, p = 0.9234; significant main effect of reward, F(1,32) = 6.76, p = 0.014). During the Preshift phase, mice entered the large-reward arm more rapidly (Fig. 1G; n = 9 mice; two-way ANOVA, significant interaction between phase and reward, F(1,354) = 6.67, p = 0.0102; posthoc pairwise comparison between large reward and small reward for the pre-shift trials, p = 1.92 e-5) and took shorter paths to the large-reward arm compared to the small-reward arm (Fig. 1H; n = 9 mice; two-way ANOVA, significant interaction between phase and reward, F(1,354) = 5.77, p = 0.0168; posthoc pairwise comparison between large reward and small reward for the pre-shift trials, p = 5.77 e-4). This pattern of fast, direct trajectories to the large reward arm is consistent with the established preference for that location before the downshift occurred. Following the unexpected reward downshift, this behavioral asymmetry was significantly reduced. The latency to enter the arm associated with the downshifted reward increased significantly after the downshift (Fig 1G; posthoc pairwise comparison between pre-shift and post-shift for large/shifted arm, p = 1.2 e-4) while the latency to enter the stable reward arm remained unchanged (Fig. 1G; posthoc pairwise comparison between pre-shift and post-shift for the small/stable arm, p = 0.93). Additionally, mice took more indirect paths toward the downshifted reward arm after the downshift (Fig 1H; posthoc pairwise comparison between pre-shift and post-shift for large/shifted arm, p = 0.0036) but showed no change in trajectories directed toward the stable reward arm (Fig. 1H; posthoc pairwise comparison between pre-shift and post-shift for the small/stable arm, p > 0.999). Together, these findings indicate that mice rapidly detected the reduction in reward value and initiated a behavioral change before an adjustment in choice preference occurred.

### Population dorsal CA1 activity at reward locations encodes reward magnitude and updates following unexpected reward downshift

The dorsal hippocampus encodes reward-related spatial memories within cognitive maps that often overrepresent reward locations ^21^ ^15^ ^22^. We asked whether CA1 also encodes specific reward magnitudes. C57BL/6 mice expressing virally injected GCaMP7f in CA1 neurons were implanted with a miniaturized 1-photon microscope (Fig. 2 A–D). Somatic Ca²⁺ activity was recorded from 1,770 – 2,364 neurons per session (134 – 486 per mouse; Extended Data Fig. 2) in 9 animals (n = 4 female, 5 male). We first examined whether hippocampal CA1 activity differentially represented the two reward magnitudes and whether these representations changed following the large reward downshift. For each neuron, Ca²⁺ event rate maps were created by calculating the normalized event rate across the linearized figure-8 maze during running (Fig. 2F, Extended Data Fig. 3). Only data recorded during forced trials were used to ensure equal sampling of the large and small reward locations. We then compared the averaged normalized event rates during running (speeds > 2 cm/s) on the large/shifted and small/stable arms for each of the sessions of interest (Fig. 2G-J). During the session when the preference criterion was met, Ca²⁺ event rates were significantly higher at the large reward location than at the small reward location (Fig. 2G, Extended Data Fig. 3; n = 9 mice, 2018 neurons; generalized linear mixed model, significant main effect of reward, t(4032) = 2.94, p = 0.003). Before the magnitude of the large reward was decreased during the Downshift session, this difference was again significant, with higher rates at the large reward location (Fig. 2H, S3; n = 9 mice, 1770 neurons; generalized linear mixed model, significant interaction between task phase and reward, t(7076) = 2.92, p = 0.003; post-hoc pairwise comparison between large reward and small reward for preshift trials p = 0.0008). Strikingly, immediately following the downshift, the difference in event rates between the previously large (downshifted) and small reward locations disappeared (Fig. 2H, S3; n = 9 mice, 1770 neurons; post-hoc pairwise comparison between large reward and small reward for postshift trials p = 0.3022). Interestingly, the normalized calcium activity did not appear to reflect reward magnitude alone; during the session where behavioral adjustment was observed, Ca²⁺ event rates were significantly lower at the shifted reward location than at the small reward location (Fig. 2I, S3; n = 9 mice, 2364 neurons; generalized linear mixed model, significant main effect of reward, t(4726) = 4.32, p = 1.63 e-5). This reduction is remarkable because both locations provided the same reward magnitude at that point in the task. To ensure that the reward magnitude representations were not influenced by ambiguous locomotor states or activity during reward consumption, data from the Downshift session were re-analyzed using a more conservative threshold to define locomotion states (5 cm/s cutoff). Normalized Ca^2+^ event rates were higher at the large reward compared to the small reward during the preshift trials, but not the postshift trials (Extended Data Fig.4; generalized linear mixed model, generalized linear mixed model, significant interaction between task phase and reward, t(7032) = 2.09, p = 0.036; post-hoc pairwise comparison between large reward and small reward for preshift trials p = 0.004; post-hoc pairwise comparison between large and small reward for postshift trials p = 0.126). The preservation of the population code at this higher speed threshold confirms that the magnitude code is a feature of active locomotion and is not driven by immobility-related transients. Notably, we observed differences in population activity between the two reward locations well before mice enter reward zones, in line with an anticipatory code for reward magnitude. Finally, we found that removing the shifted reward entirely also resulted in lower Ca^2+^ event rates compared to the intact small reward (Figure 2J, S3; n = 8 mice, 1939 neurons; generalized linear mixed model, significant main effect of reward, t(3874) = 6.00, p = 2.065 e-9). These results indicate that CA1 representations of reward magnitude are flexible and update according to changes in magnitude and behavioral preference.

**Figure 2.**
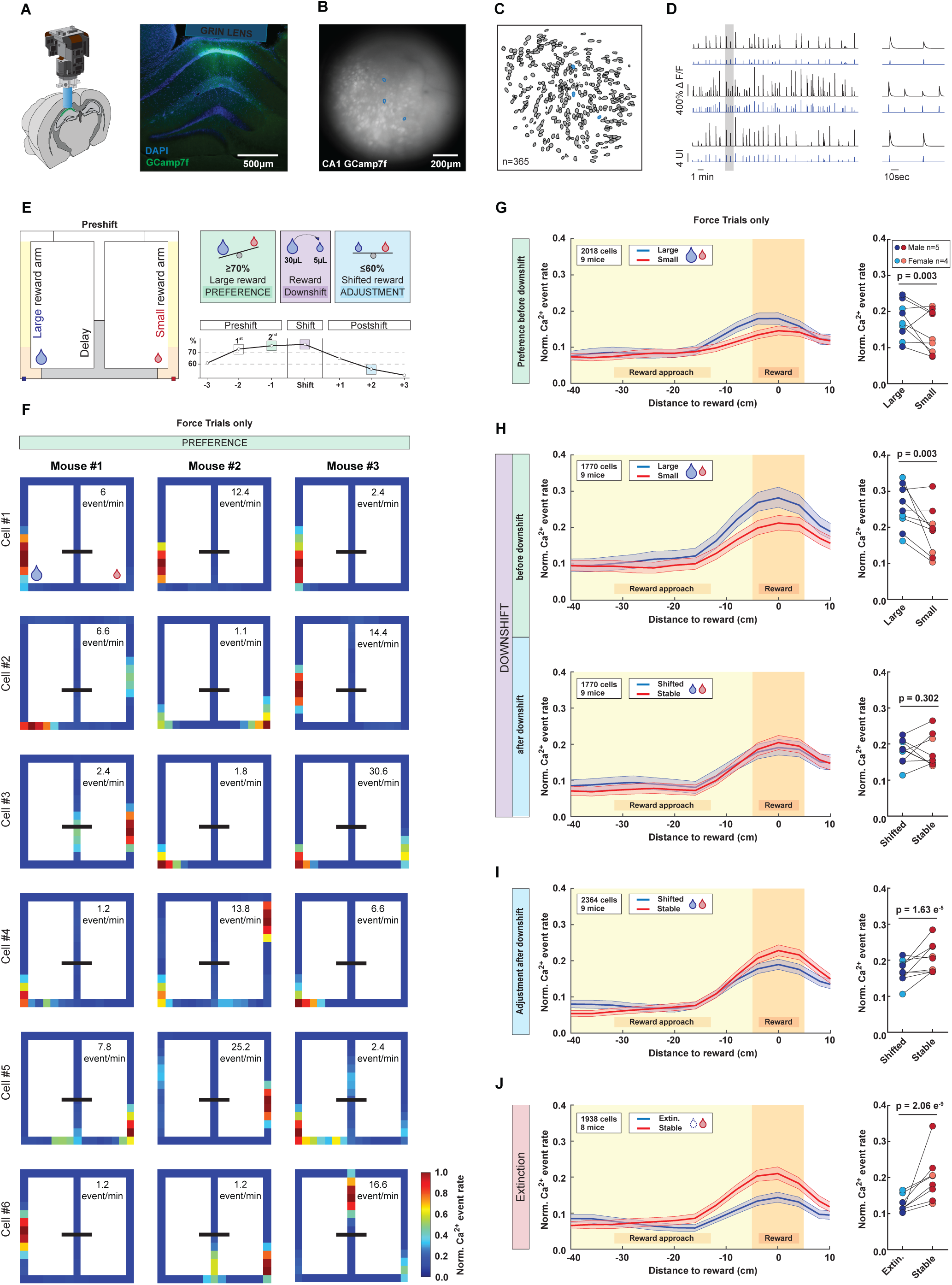
Population code for reward magnitude in dorsal CA1. (A) *In vivo* freely moving Ca²+ imaging of dorsal CA1 hippocampal activity during the reward devaluation task. Left: Schematic of UCLA Miniscope v4 with GRIN lens implant targeting dorsal CA1. Middle: Histological section showing GRIN lens placement above the stratum oriens of CA1, with AAV-syn-jGCaMP7f expression (green) and nuclear staining (DAPI, blue). Scale bar, 500 µm. (B) Maximum intensity projection of a 5-minute calcium imaging session from dorsal CA1. Scale bar, 500 µm. (C) Extracted spatial footprints using a sparse nonnegative matrix factorization (NMF) approach implemented in Minian. (D) Representative calcium traces from three example cells, with matched cell identities shown on the footprints spatial map (blue), raw calcium fluorescence traces (black), and deconvolved activity signals (blue). (E) Left: Schematic of the figure-eight maze, highlighting the key locations: delay zone (grey), stem arm (white), reward arms (yellow), and reward zone (orange). Right: Timeline of the reward devaluation task, showing the different phases: preshift phase (light green), shift phase (light purple), and postshift phase (light blue). (F) Examples of calcium event activity maps from three animals during the last day of preshift phase (forced choice trials), when the mice developed a preference for the large reward. The maps represent calcium event rates, with colors ranging from blue (no activity) to red (highest event rate). Each bin of the linearized map has a size of 4 cm. Black lines indicate the delay zone door. The locations of the large and small rewards are shown. Calcium event rates (event/min) are provided for each cell. (G-J) Trajectory-dependent neuronal activity during Preshift, Downshift, and Postshift sessions. (Left) The binned normalized calcium event rate during running for the reward arm (“Reward approach”) and reward zone (“Reward”) during forced choice trials, pooled across all neurons from all mice (n = 9). Blue traces correspond to turns toward the large/shifted reward, and red traces correspond to turns toward the small/stable reward. Shaded areas represent 95% confidence intervals. (Right) Mean normalized calcium event rate at reward locations averaged across neurons for each mouse across phases of the reward downshift task. Male mice are shown in dark blue and dark red, female mice are shown in light blue and light red. P-values shown for generalized linear mixed effects model comparing activity across rewards.

### Reward population codes are not driven by overrepresentation of CA1 place cells at large reward

We next asked whether the varied Ca²⁺ event rates observed at the reward locations were driven by differences in place cell activity. We identified place fields of CA1 neurons with stable spatial activity from two-dimensional ratemaps of Ca²⁺ events (Fig. 3A). We found that CA1 place cells represented activity across the entirety of the figure-8 maze (Fig. 3A, B, n = 1628 place fields). Consistent with previous reports, reward locations showed a higher density, or overrepresentation, of place fields relative to the rest of the maze. There was a similar percentage of fields at the large and small reward locations prior to reward downshift (Figure 3C; Chi squared test, *χ*2(1,1627) = 1.94, p = 0.164). Furthermore, quantitative comparisons of place cell properties revealed no significant differences between the fields located at the large (n = 326) and small (n = 297) reward locations; there was no difference in peak Ca²⁺ event rate (Fig. 3C, Wilcoxon rank sum test, z = −1.25, p = 0.208), mean in-field Ca²⁺ event rate (Fig 3C; Wilcoxon rank sum test, z = −1.30, p = 0.191), or place field size (Fig 3C; Wilcoxon rank sum test, z = 0.71, p = 0.475). These results indicate that reward population codes are not driven by overrepresentation of, or higher rate, CA1 place cells at the large reward location.

**Figure 3:**
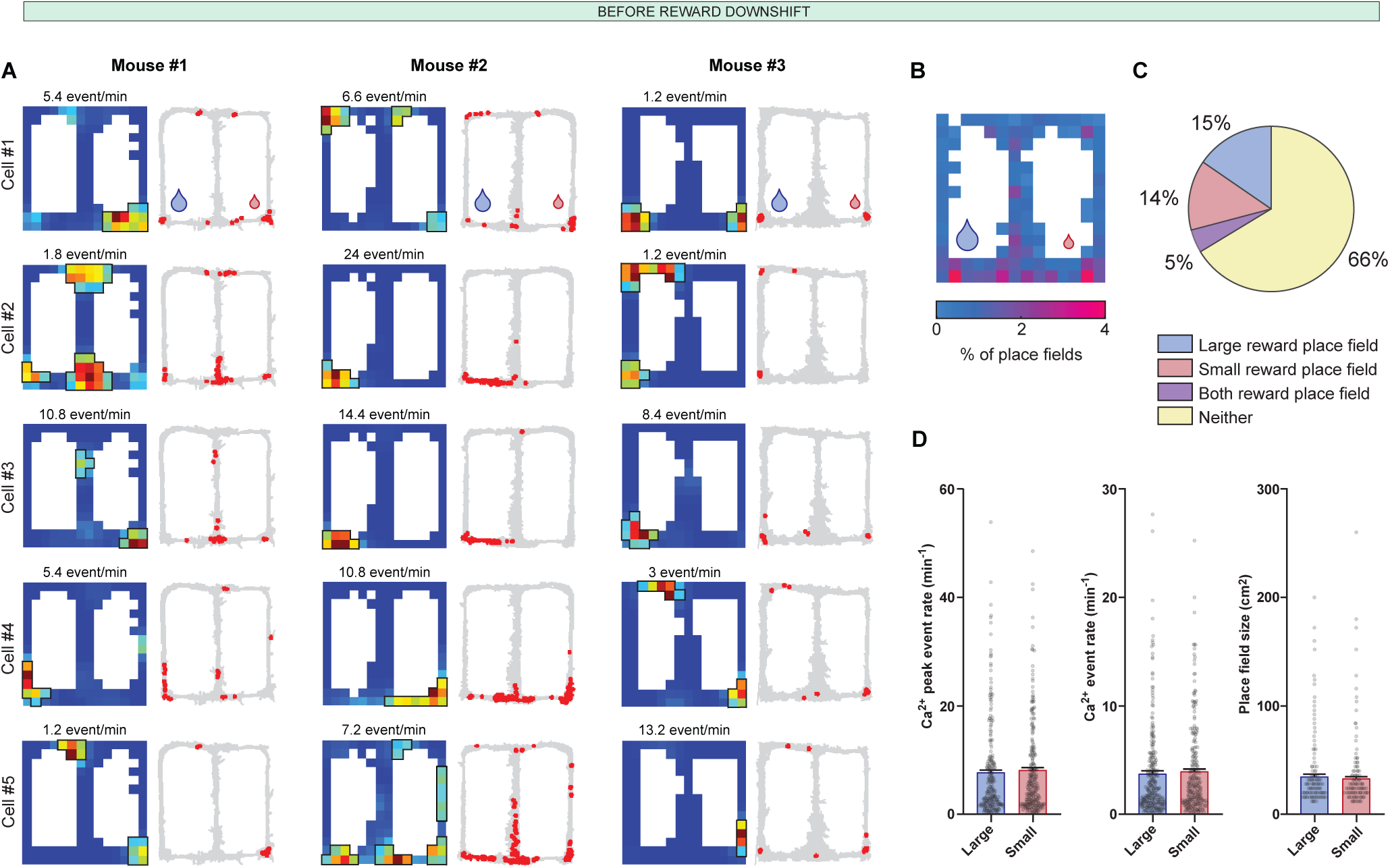
Representation of reward magnitudes is not driven by spatial encoding of place cells with place fields at reward locations. (A) Representative place cells activity maps with place fields at reward locations prior to reward magnitude downshift and corresponding locomotion trajectory maps showing the individual calcium events (red dots) from each cell. (B) The distribution across the maze of the peak firing locations for each place field before the shift on the Downshift day. Reward locations are overrepresented relative to the rest of the maze. (C) There was no difference between the proportion of place fields located proximal to the large reward compared to the small reward. P-value shown for Chi-squared test comparing proportion of large and small rewards. (D) There was no difference in the peak or average in-field firing rates of place cells located within 5 cm of the large or small reward. P-value shown for Wilcoxon rank sum test comparing values at large and small rewards. Error bars indicate SEM.

### Large reward-tethered neurons are the primary drivers of hippocampal reward-magnitude coding

To gain further insight of what cell types were contributing to the population reward magnitude code, we identified neurons that had increased activity at the large reward prior to reward downshift using a criterion that was much less stringent than place field identification (i.e., a firing rate at the large reward that was one standard deviation above baseline across the maze; Fig. 4A; n = 479 place cells). As expected, approximately half of these cells had place fields located at the large reward or place fields located at both rewards (Fig. 4B). The next largest population consisted of neurons that lacked a place field at the large reward and had a place field located elsewhere on the maze (Fig. 4A-C). We termed these cells ‘large reward-tethered’ (LRT) place cells because their map-wide activity pattern is tied to moderate activity at the large reward location, even though they also represent spatial regions distant from the reward on the maze.

**Figure 4:**
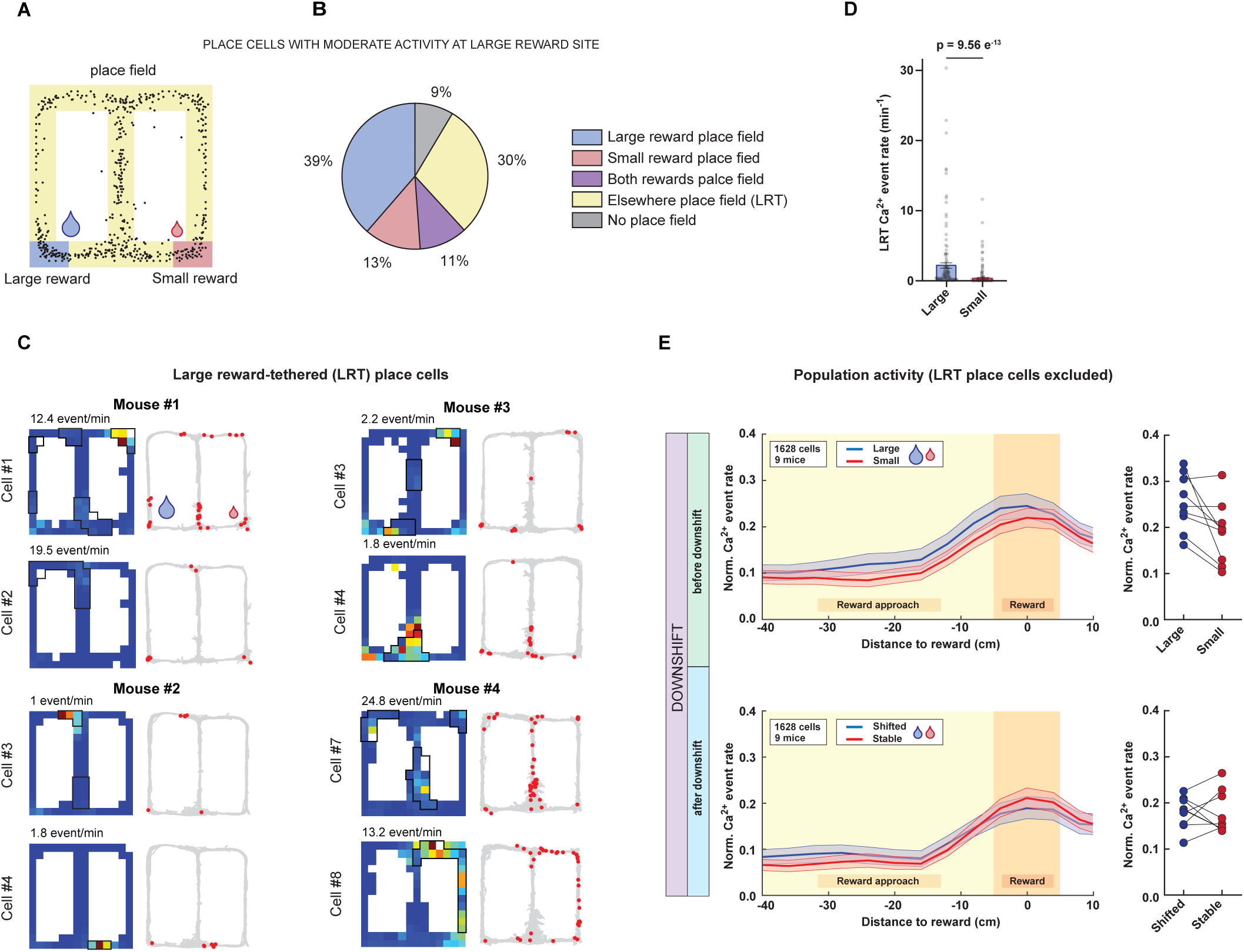
Large reward-tethered (LRT) place cells provide the cellular substrate for reward-magnitude coding. (A) Spatial distribution of place field centers of mass for all neurons that have a firing rate of at least 1 standard deviation above the mean at the large reward for the preshift forced trials during the Downshift day. The large reward zone shown in blue, small reward zone shown in red, other locations in yellow. (B) Pie chart shows the identities of cells that have a firing rate of at least 1 standard deviation above the mean at the large reward for the pre-shift forced trials for the downshift day. (C) Activity maps of cells exhibiting moderate activity at the large reward. Corresponding locomotion trajectories show individual calcium events (red dots) for each cell. Peak firing rate is shown in the inset for each map. (D) Column plots represent calcium event rate at the large (blue) and small (red) reward location for LRT place cells. Error bars indicate SEM. P-value shown for Wilcoxon signed rank test comparing values at large and small rewards. (E) Trajectory-dependent neuronal activity during Preshift sessions of non LRT place cells. (Left) The binned normalized calcium event rate during running for the reward arm (“Reward approach”) and reward zone (“Reward”) during forced choice trials, pooled across all neurons from all mice (n = 9). Blue traces correspond to turns toward the large reward, and red traces correspond to turns toward the small reward. Shaded areas represent 95% confidence intervals. (Right) Mean normalized calcium event rate at reward locations averaged across neurons for each mouse before the reward downshift.

Given the prevalence of LRT place cells, we wondered whether they contributed to the reward magnitude code. Quantifying the firing rates of LRT place cells revealed a significantly higher discharge rate at the large-reward location compared to the small-reward location (Fig. 4D; Wilcoxon signed rank test, z = 7.14, p = 9.56 e-13). We then asked whether the population activity difference in event rates observed at the unequal reward locations (Fig. 2H) was primarily driven by the LRT place cell sub-population. To test this, we removed LRT place cells and re-analyzed the data (Fig. 4E). Remarkably, the exclusion of this single sub-population completely abolished the magnitude-dependent divergence in population activity across task phases; without LRT place cells, there was no significant difference in event rates between the large and small reward locations during either the preshift or postshift trials (Fig. 4E; generalized linear mixed model, no interaction between task phase and reward, t(6508) = −1.80, p = 0.0726; no main effect of reward, t(6508) = −0.06, p = 0.950). These results demonstrate that the hippocampal representation of reward magnitude is critically dependent on the recruitment of LRT place cells, which suggests that this cell type provides a specialized cellular substrate for encoding reward magnitude within the dorsal CA1 map.

### Preferential remapping of LRT place cells following reward-contingency updates

Previous work shows that extinction of rewards can destabilize spatial representations of the entire environment, indicating that the hippocampus builds new maps for new reward contingencies ^16^. We wondered if downshifting would have a similar impact. In other words, is a reduction in reward magnitude sufficient to induce remapping throughout the entire environment?

We were especially interested in whether LRT place cells and non-LRT place cells would behave similarly after downshifting. In order to quantify the change in firing fields, we calculated the spatial correlation values between the preshift and postshift firing rate maps across LRT and non-LRT place cells (Fig. 5A). We excluded the reward zones in order to capture potential remapping not related to the moderate firing at the large reward location. We found that the spatial stability was lower in LRT place cells (n = 142) compared to non-LRT place cells (n = 681) (Fig 5B; Wilcoxon rank sum test, z = −4.33, p = 1.49 e-5). To visualize the extent of this remapping across the maze, we then constructed population maps (Fig. 5C) of the figure-8 maze before and after the reward downshift. Population vector correlations calculated with LRT place cells were qualitatively lower than for non-LRT place cells (Fig. 5D). Further, these lower population vector correlation values were seen across the entire extent of the maze indicating that LRT cells changed their firing fields following reward downshift. Overall, these changes indicate that reward-responsive neurons with place fields outside of the reward locations undergo selective reorganization following a reward downshift. Rather than reflecting a global disruption of spatial coding across all neurons, this effect was specific to neurons engaged at the large-reward location. This suggests a context-dependent modulation of spatial activity patterns.

**Figure 5:**
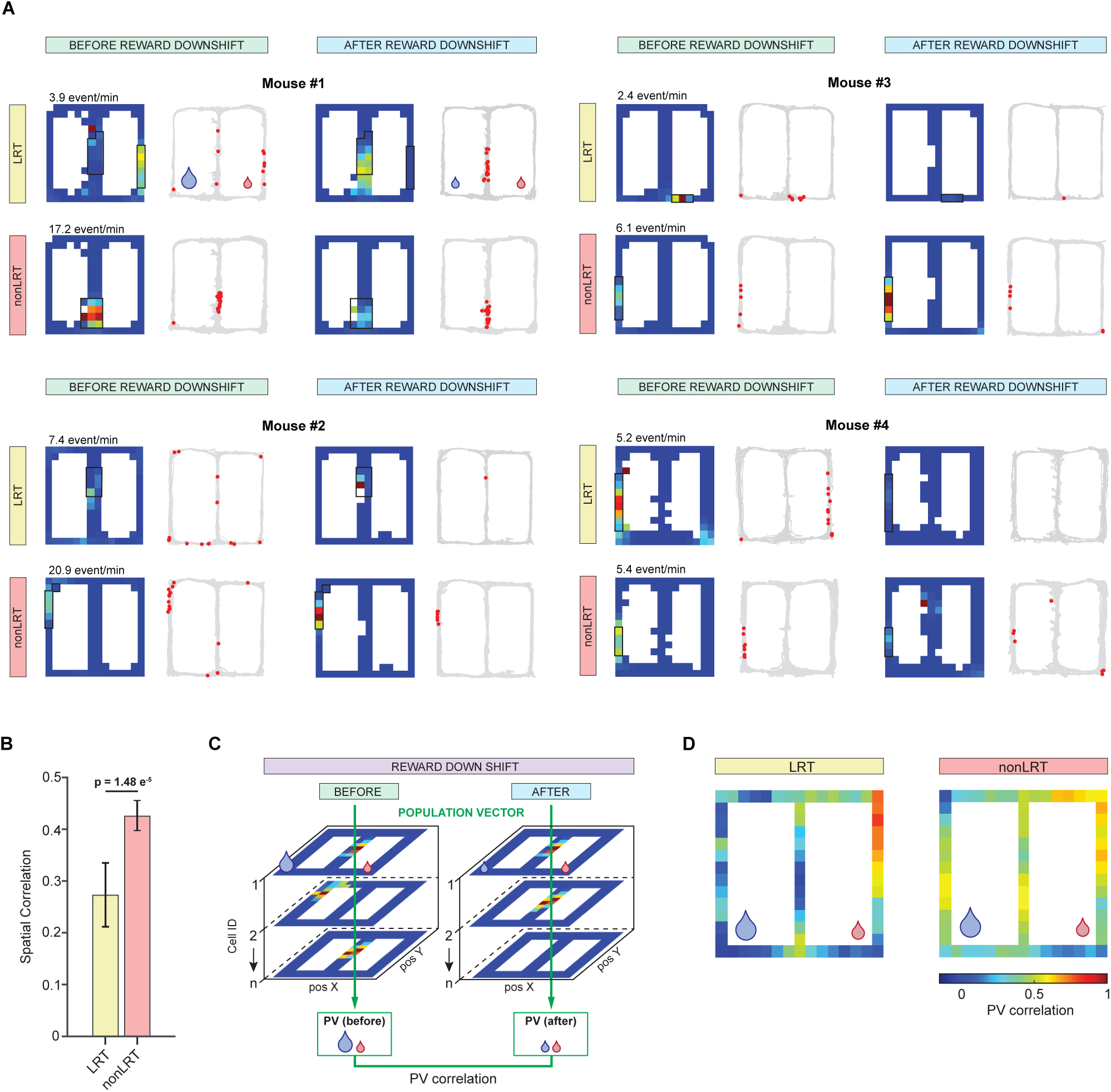
Selective remapping of Large reward-tethered (LRT) place cells following reward magnitude downshift. (A) Activity maps of place cells with (LRT) and without (non-LRT) reward-associated activity at the large-reward location prior to downshift. Corresponding locomotion trajectories show individual calcium events (red dots) for each cell. Peak firing rate is shown in the inset for each map. (B) Spatial correlation values calculated using maps before and after the downshift (excluding reward zones) were lower for LRT place cells compared to non-LRT place cells. P-value shown for Wilcoxon rank sum test comparing values at LRT and non-LRT place cells. Error bars indicate standard error of the mean. (C) Schematic representation of a population vector correlation representing the changes of combined calcium events activity of a neural ensemble across the maze. (D) Population of LRT place cells exhibit lower population vector correlations across the maze on the Downshift day compared to non-LRT place cells.

## DISCUSSION

In this study, we investigated how the dorsal hippocampus integrates reward values into the cognitive map and how these representations reorganize when expectations are violated. Our results demonstrate that CA1 population activity encodes reward value through elevated event rates at high-value locations. Further, CA1 activity appears to reflect expectation-dependent value, rather than absolute magnitude. Concomitant with the preference adjustment after reward downshift, activity at the shifted location fell significantly below that of the stable small-reward location, despite identical reward sizes at both locations. This implies that the hippocampus is doing more than integrating a sensory mismatch; it is actively re-evaluating value. The magnitude suppression might represent a distinct neural signature of the internal model update or negative incentive contrast ^12,23^, suggesting that the hippocampus acts as an active comparator that evaluates the mismatch between “what is expected” and “what is received” to drive a shift in the cognitive map ^7–9^. Crucially, this departure from a pure magnitude code could suggest that CA1 activity reflects an internal strategy where the ensemble prioritizes the stable, consistent reward over the downshifted one, effectively de-valuing the previously large reward to guide future choice.

Our results further show that the hippocampal value code is supported by a distinct functional class of neurons, which we term large reward-tethered (LRT) place cells. LRT place cells multiplex spatial and value related information by exhibiting selective moderate firing at the large reward location while maintaining place fields elsewhere. This finding aligns with the emerging view that CA1 contains multiple representations differentially coupled to static environmental features versus dynamic, task-relevant variables ^24,25^ ^15,17,26^. By maintaining distal place fields and selective reward responses, LRT place cells may act as primary anchors within the cognitive map, linking environmental context to expected outcome value. Instead of providing a static neural scaffold, these neurons likely facilitate the flexible magnitude coding observed at the population level, as their selective moderate activation at reward locations may prime them for plasticity when expectations are violated. Therefore, while value linked neurons are preferentially updated, non-LRT populations preserve a more stable spatial scaffold ^17,27,28^.

Following an unexpected downshift in reward magnitude, LRT place cells undergo rapid remapping that precedes behavioral adjustment of reward location preference. This suggests that hippocampal neuronal dynamics could be one of the drivers in the updating of internal value representations rather than merely reflecting altered behavioral strategies. The selective remapping of LRT place cells points to an interaction between spatially modulated neuronal activity and neuromodulatory gating. While the hippocampal–ventral tegmental area (VTA) loop provides a plausible substrate for reward prediction errors ^29^, the rapid remapping of LRT place cells may be further gated by a parallel dopaminergic system reflecting changes in predicted outcomes ^30^. Specifically, the locus coeruleus (LC) co-releases dopamine into the dorsal hippocampus, providing a high-potency signal for environmental novelty that enhances synaptic plasticity in an activity-dependent manner ^31^ ^32,33^. Future studies manipulating LC or VTA projections to CA1 during the downshift window will be essential to confirm whether these neuromodulatory inputs are the causal gatekeepers of LRT remapping.

Our results raise a fundamental question of information flow: is the hippocampus the primary detector of contingency shifts during goal directed navigation? While simple reward seeking can eventually become hippocampal independent ^34^, rapid adaptation to downshifted outcomes requires a high fidelity comparison of current sensory input against stored spatial expectations ^9,35^. Our previous work established the necessity of the dorsal hippocampus for behavioral flexibility, as animals cannot adjust anticipatory behavior following a reward downshift without an intact hippocampus ^20^. Our current physiological results extend this finding, suggesting that the hippocampus is effectively a ‘first detector’ – exhibiting remapping that precedes behavioral adjustment. In this framework, LRT place cells may be critical for signaling the initial invalidation of expected reward value, a role supported by recent evidence showing that midbrain dopamine responses to expectation violations are hippocampal dependent ^36^. By broadcasting this updated value signal to downstream regions like the nucleus accumbens and VTA ^37,38^, the hippocampus may initiate a brain wide circuit update and facilitate new behavioral strategies.

By coupling the selective remapping of LRT place cells with the stable spatial representation of non-LRT place cells during reward downshift, the hippocampal circuit effectively balances environmental continuity with adaptive flexibility. Failure of this process may underlie the perseverative, maladaptive behaviors observed in depression, anhedonia, and substance use disorders ^39–41^. If LRT-like neurons remain pathologically coupled to outdated rewards, they may bias behavior towards inefficient strategies despite changes in outcome value. By identifying a population-level mechanism that links expectation, value, and spatial context, we provide a framework for investigating how flexible behavior is implemented, and disrupted, at the level of defined neural circuits.

## METHODS

### Animals

Adult C57BL/6 mice were used across all experiments. For the c-Fos quantification experiment, a cohort of 22 C57BL/6J mice (12 males, 10 females; 11–12 weeks old) was used at Mount Holyoke College Massachusetts (MHC). Mice were housed individually under a reversed 12:12-hour light/dark cycle in a climate-controlled environment (22 ± 2 °C; 65% humidity). All behavioral experiments were performed during the dark cycle. All procedures for this experiment were approved by the Mount Holyoke College Institutional Animal Care and Use Committee. All other experiments were done at the University of California, Irvine and approved by the Institutional Animal Care and Use Committee. 10 mice were used (5 males, 5 females; 12 weeks old). Mice were maintained under housing conditions identical to those described above. Animals were handled daily for three days prior to the start of behavioral testing. Body weight was recorded prior to the start of the experiment and defined as 100% (*ad libitum* baseline). Citric acid (2%) was added to the drinking water to facilitate voluntary water restriction and enhance motivation for sucrose rewards (5% sucrose solution). While on this regiment, mice maintained at least 85% of their baseline body weight.

### Automated Reward Devaluation Task

The task was performed using a figure-8 maze (56 cm x 56 cm, with 5 cm wide pathways), equipped with five automated doors controlled by an Arduino interface that guided the mouse’s movement in a single direction as indicated by the arrows in Figure 1A. The opening and closing of the doors were triggered based on the animal’s position, which was continuously monitored by a camera (TECKNET 1080p). A custom Bonsai Rx81 workflow was implemented to control the automated behavioral task and to acquire, synchronize, and process multiple data streams in real time. The two corners of the maze designated as the reward areas were fitted with automated ports (Bpod port, Sanworks), which provided visual cues and delivered liquid sucrose rewards. The reward devaluation task consists of six phases: Habituation, Preshift training, Preshift testing, Shift testing, Postshift testing, and Extinction.

#### Habituation

The habituation phase consisted of two days, each consisting of a 20 minute exploratory session. On the first day mice received unlimited rewards (5 μL 5% sucrose water) at both reward ports. On the second day animals were required to nose poke to break the infrared beam inside the port to receive rewards.

#### Preshift training

Animals learned to associate each reward port with a distinct reward magnitude. One port was baited with 5 µL of a 5% sucrose solution, while the other was baited with 30 µL of 5% sucrose solution. Trials (n = 40) were forced and presented in a pseudorandom order, with an equal distribution of left and right trials. Between trials mice were confined to the delay zone for an inter-trial interval (ITI) of 15 seconds. This phase lasted until mice visited each port at least 10 times (2-3 days).

#### Preshift testing

During this phase, animals continued learning reward associations while their preferences were assessed to evaluate the learning outcome. The first 20 trials of each session were free choice and mice could freely choose either reward port to receive the associated reward (5 µL or 30 µL). Beginning with trial 21, forced trials were introduced, consisting of 10 left and 10 right trials presented in a pseudorandom order. Mice progressed to the next phase only after demonstrating a strong preference (≥70%) for the larger reward location on 2 out of 3 consecutive days.

#### Shift testing

The shift day consisted of 60 trials in total. The session began with 20 preshift forced trials, during which mice received the previously established rewards: 30 µL at the large reward location and 5 µL at the small reward location. In the subsequent 20 free choice trials (trials 21-40), the large reward was downshifted to match the smaller reward, with both reward locations delivering 5 µL. Trials 41-60 consisted of forced shift trials during which mice received 5 µL of reward at both reward locations.

#### Postshift testing

All Postshift testing sessions consisted of 20 free choice trials followed by 20 forced trials (pseudorandomly ordered with 10 left and 10 right trials). In all trials, mice received 5 µL of a 5% sucrose solution at both reward locations. Postshift testing continued daily until mice demonstrated behavioral adjustment to the reward downshift, indicated by a preference of 60% or less for the shifted reward.

#### Extinction

During the extinction phase, no reward was provided on the previously downshifted location. Mice ran 20 free choice trials followed by 20 pseudorandomly presented forced choice trials. This phase lasted for two days.

### Non-automated reward devaluation

c-Fos experiments were conducted in a modified figure-8 maze using a reward downshift paradigm adapted from the main task. In contrast to the liquid sucrose rewards used in the primary experiments, rewards consisted of sugar pellets (20 mg each). This modification allowed us to preserve the overall reward contingencies and behavioral structure while accommodating equipment differences between laboratories. Mice were maintained at 85–90% of their ad libitum body weight through food restriction starting three days prior to the study. Water was provided ad libitum throughout the experiment. Following three days of handling and a 15-minute habituation to the maze, mice underwent a multi-stage behavioral protocol. The Preshift training phase lasted 7 days (Sessions 1–7); each session consisted of 6 trials, beginning with 2 free-choice trials followed by 4 forced-choice trials (2 left, 2 right, counterbalanced in order) to ensure equal sampling of both reward magnitudes. During the subsequent Preshift testing phase (Sessions 8–10), which lasted 3 days, mice performed 6 free-choice trials per session to assess reward discrimination between the large reward (6 pellets) and the small reward (1 pellet). By session 10, all mice reached a stable preference criterion of ≥70% for the large reward. On the Reward downshift day (Session 11), the large reward was unexpectedly reduced to 1 pellet to match the small reward magnitude. The downshift session consisted of 6 free-choice trials to evaluate the behavioral adjustment to the incentive devaluation. To capture the peak of immediate early gene expression, mice were perfused 90 minutes after the conclusion of the downshift session.

### c-Fos Quantification

#### Histology and Immunohistochemistry

Mice were sacrificed 90 minutes after the final behavioral session on day 11 to capture peak c-Fos expression. Subjects were anesthetized with sodium pentobarbital at a dose of 80 milligrams per kilogram and transcardially perfused with 0.01 molar phosphate-buffered saline, followed by 4% paraformaldehyde in phosphate-buffered saline. Brains were extracted and post-fixed in 4% paraformaldehyde for 24 hours, then transferred to a 30% sucrose solution for at least 48 hours. Coronal sections with a thickness of 20 µm were obtained using a ThermoScientific Cryostar NX50 cryostat. Every third section was processed for free-floating c-Fos immunohistochemistry to provide systematic sampling. This sampling began at the level of the anterior commissure and continued posteriorly through the entire length of the dorsal hippocampus. Sections were incubated in 1% hydrogen peroxide to block endogenous peroxidases and then placed in a blocking solution of 10 percent normal donkey serum for 30 minutes. Tissue was subsequently incubated for 24 hours at 4 degrees Celsius in a primary rabbit anti-c-Fos antibody from Millipore at a dilution of 1 to 5000 in phosphate-buffered saline containing 0.8% Triton X-100 and 0.1% bovine serum albumin. Following the primary incubation, a biotinylated donkey anti-rabbit secondary antibody from Jackson ImmunoResearch was applied at a 1:500 dilution for 90 minutes. Signal amplification was achieved using a Vector Elite ABC kit at a 1:200 dilution, and the signal was visualized with a Vector SG HRP chromogen kit for 7 minutes. Sections were counterstained with nuclear red, dehydrated through a series of graded ethanols from 50 to 100%, cleared in xylene, and coverslipped using Permount.

#### Image Acquisition and Quantification

Brightfield images of the dorsal hippocampus, including the CA1, CA3, and dentate gyrus, were acquired using a standard light microscope with consistent exposure settings. Automated cell counting was performed using QuPath software version 0.6.0. Regions of interest were manually annotated for hippocampal subfields, and c-Fos-positive nuclei were identified using the cell detection tool based on optical density sum. Detections were manually verified by two blinded experimenters. Neuronal density was calculated as cells per square millimeter (mm²) by dividing the total number of c-Fos-positive cells by the area of the region of interest in mm².

### Surgical Procedure for Calcium Imaging

Each mouse underwent two surgeries: (1) stereotaxic viral injection for calcium indicator expression in dorsal CA1, and (2) cranial window implantation for GRIN lens placement.

#### Viral Expression of Genetically Encoded Calcium Indicator

To enable in vivo calcium imaging in dorsal hippocampal CA1, we injected an adeno-associated virus encoding GCaMP7f (pGP-AAV-syn-jGCaMP7f-WPRE; Addgene; 1:4 dilution in 0.9% NaCl, final volume 600 nL per mouse; ≥10¹³ vg/mL titer). Prior to surgery, mice were weighed to calculate doses for lidocaine and carprofen. Anesthesia was induced with isoflurane (3–4%) in a 25% O₂ / 75% air mixture and maintained at 1–2% during surgery. Mice were placed in a stereotaxic frame (David Kopf Instruments). Subcutaneous carprofen (5 mg/kg; 0.1 mL/10 g body weight) was administered pre-operatively. Body temperature was maintained at 37–38°C using air-activated heat packs (HotHands), and ophthalmic ointment (Soothe) was applied to prevent corneal drying. After confirming the surgical plane via absence of reflexes, fur was removed and the scalp disinfected with 10% povidone-iodine. A local anesthetic (lidocaine 2%; 4 mg/kg, 0.008 mL/10 g body weight, s.c.) was administered before a midline incision. Coordinates for virus injection (−1.9 mm AP, −1.6 mm ML relative to bregma) were marked. Burr hole was drilled, and the dura removed. Virus was pressure-injected at two sites (−1.5 and −1.6 mm ML; DV −1.1 mm from dura) using a glass pipette and microinjection system (MO-10, Narishige). Each site received 300 nL over 3 min, with 1-minute pre- and post-injection dwell times to reduce backflow. Following injection, the scalp was sutured and treated with antibiotic ointment (Neosporin). Mice were monitored during recovery and housed individually for two weeks prior to lens implantation.

#### Cranial Window Implantation for CA1 Imaging

Two weeks after viral injection, mice underwent implantation of a gradient refractive index (GRIN) lens above dorsal CA1. Baytril Placebo (5 mg tablet, Bio-Serv) was administered orally 48 h before surgery, and replaced with antibiotic-containing Baytril (0.5 mg, Bio-Serv) 24 h pre-operatively. On the day of surgery, mice were weighed and given buprenorphine (0.1 mg/kg, s.c.) and dexamethasone (4 mg/kg, i.p.). lidocaine (2%; 4 mg/kg, s.c.). Mice were anesthetized and positioned as previously described. After confirming anesthesia depth, mice were injected with lidocaine (2%; 4 mg/kg, s.c.), the scalp was removed and the skull cleaned with 3% hydrogen peroxide for 40 s. The bone surface was scored to improve adhesion, then coated with Optibond (Kerr) and cured under UV light (Dentmate Ledex; 3 × 30 s). A 2-mm diameter craniotomy was made at the prior injection site. Throughout the procedure, the cortex was kept moist with sterile saline (0.9%) delivered via blunt cannula. Cortical tissue was aspirated with a 27G needle under vacuum until the alveus was exposed. A 1-mm GRIN lens (Gofoton) was lowered to the alveus and fixed in place with dental cement (Tetric EvoFlow). The exposed lens was sealed with silicone elastomer (Kwik-Sil, WPI) to prevent damage before miniscope base-plating. Mice received post-operative carprofen (5 mg/kg, s.c.) and were allowed to recover under heat. They were monitored daily for three days before head-plate attachment.

#### Miniscope Base-Plating

Three to five days post-implantation, base plates were affixed for miniscope imaging. Mice were anesthetized and placed in a stereotaxic frame with a mounted UCLA Miniscope V4. The silicone cap was removed and the GRIN lens cleaned with 70% ethanol. The miniscope was aligned with the implanted lens to visualize GCaMP fluorescence in CA1. Once optimal imaging was achieved, the aluminum base plate was secured with dental cement and the scope detached. A protective plastic cap was placed on the base plate, and black nail polish applied to all exposed cement to reduce ambient light interference. Mice recovered under heat and were monitored post-operatively.

### Calcium Imaging Data Processing

Calcium imaging data were processed using the open-source pipeline Minian to extract spatial footprints, fluorescence traces, and corresponding deconvolved activity. To identify baseline signal drift, each recording was segmented into one-minute intervals, and local minima (valleys) were analyzed. A cell was flagged as exhibiting drift if any valley exceeded 0.1 arbitrary units (A.U.) for a continuous period of at least five minutes. These automatically detected drift events were then manually reviewed using a custom graphical user interface (GUI), which displayed both the raw calcium trace and deconvolved signal for verification. Cells identified as drifting were excluded from all subsequent analyses. Additionally, cells with fewer than four detected calcium events during the 30-minute recording session were removed. For the remaining population, calcium event times were extracted by peak detection applied to the deconvolved traces.

#### Position and speed tracking

Animal locomotion activity was recorded using a 1080p webcam (TECKNET 1080p Webcam) at a frame rate of 30 fps. Online position estimation was performed in Bonsai RX by computing the center of mass of the mouse. Running speed was derived from body position and smoothed Gaussian kernel (σ = 500 ms).

#### Calcium event rate at reward locations

To examine hippocampal neuron firing patterns leading up to the reward locations (Fig. 2), we created one-dimensional ratemaps for each neuron. Only data from forced choice trials were included. The maze was linearized into 4 cm spatial bins. The firing rate for each cell was calculated by dividing the number of calcium events per bin by the time spent in that bin. Only times when the mouse was traveling over 2 cm/s were included. The ratemaps were then smoothed with a Gaussian kernel (σ = 6 cm). Firing rate maps were constructed separately for large reward trials and small reward trials and normalized by the max firing rate recorded for that unit that day. Cells firing less than 10 times during the entire session were excluded.

#### Place cells identification

Firing rate maps were constructed for each neuron similar to our prior studies ^42^. The maze was divided into 4 cm^2^ bins and the number of spikes that occurred in each bin was divided by the time the mouse spent in the bin. Spikes and times that the mouse was moving less than 2 cm/s were excluded. The firing rate maps were then smoothed with a two-dimensional Gaussian kernel (σ = 6 cm). To test for spatial stability, maps were created for the first and second half of each phase (i.e., both halves of the forced and free choice trials). Additionally, maps were constructed for even numbered trials and odd numbered trials separately. The Pearson correlation coefficient was assessed between first half and second half maps and even and odd maps. The average of these two values was taken as a measure of spatial stability for that task phase. Calcium event times were then circularly shuffled (at least ±1 min away from the original timepoints) and correlation coefficients were recalculated 500 times to create a null distribution. For statistical analysis of spatial stability, the Fisher’s transformation of the Pearson’s correlation was used. A cell was considered a place cell if its spatial stability exceeded the 95^th^ percentile of the null distribution in at least one task phase. Cells firing less than 10 times during the entire session were excluded. Two-dimensional and one-dimensional ratemaps were constructed separately.

To identify place fields, two-dimensional rate maps constructed from the entire day were used. Potential place fields were identified as portions of the firing rate map that exceeded at least 25% of the maximum firing rate. In order to be included for further analysis, place fields had to be at least 10 cm^2^ and had to have a minimum in-field peak firing rate of at least 10% of the maximum firing rate. Place fields were considered to be near the small or large reward if any part of their field was within 5 cm of the reward port.

#### Identification of large reward-tethered (LRT) place cells

To identify cells with activity at the large reward, we examined the firing rate maps at the large reward during the pre-shift trials on the shift day. We z-scored the firing rate maps during runs to the large and small rewards. To be considered a large reward-tethered place cell, the mean of the binned firing rate at the bins adjacent to the large reward had to have a z-score of 1 or higher. Cells with identified fields at the large reward were excluded.

### Statistical Analysis

All statistical analyses were performed in MATLAB unless otherwise stated. To analyze behavior, we used a two-way ANOVA to test the effects of reward location and trial phase (*anovan* in MATLAB). Post-hoc pairwise comparisons were made using Tukey’s correction for multiple comparisons (*multcompare* in MATLAB). To analyze the normalized calcium event rates across reward locations we used a generalized linear mixed model design (*fitglme* in MATLAB). Mouse and unit nested within mouse were set as random factors, and reward identity was included as a fixed factor. When analyzing the Downshift day, task phase (i.e., trials prior to the shift and trials after the shift) was also included as a fixed factor. Posthoc pairwise comparisons used the Bonferroni correction to control for multiple comparisons. To compare the proportion of fields at the large and small rewards, a Chi-squared test was used (*crosstab* in MATLAB). To compare cFos expression, place field properties across reward locations, and spatial correlation coefficients across LRT and non-LRT cells, we used a Mann Whitney/Wilcoxon rank sum test (*ranksum* in MATLAB). To compare the firing rate of LRT place cells at the large and small reward locations, we used a Wilcoxon signed rank test (*signrank* in MATLAB).

## ACKNOWLEDGEMENTS

The following funding supported this work. R01NS128222 to L.A.E., Vorwerk Family Funds to H.A., Harap Family Funds to K.B.M.

## AUTHOR CONTRIBUTIONS

Conceptualization, L.A.E. and M.S.; data curation, N.M. and M.M.D.; formal analysis, M.M.D.; funding acquisition, L.A.E., M.S., H.A., and K.B.M., investigation, N.M., B.L.B, G.I.M, H.A., K.B.M,; methodology, N.M. and M.M.D.; project administration, L.A.E. and M.S.; supervision, L.A.E. and M.S.; visualization, = N.M., and M.M.D.; writing – original draft, reviewing and editing, N.M., M.M.D., M.S., and L.A.E.

## EXTENDED DATA

**Extended Data Figure 1:**
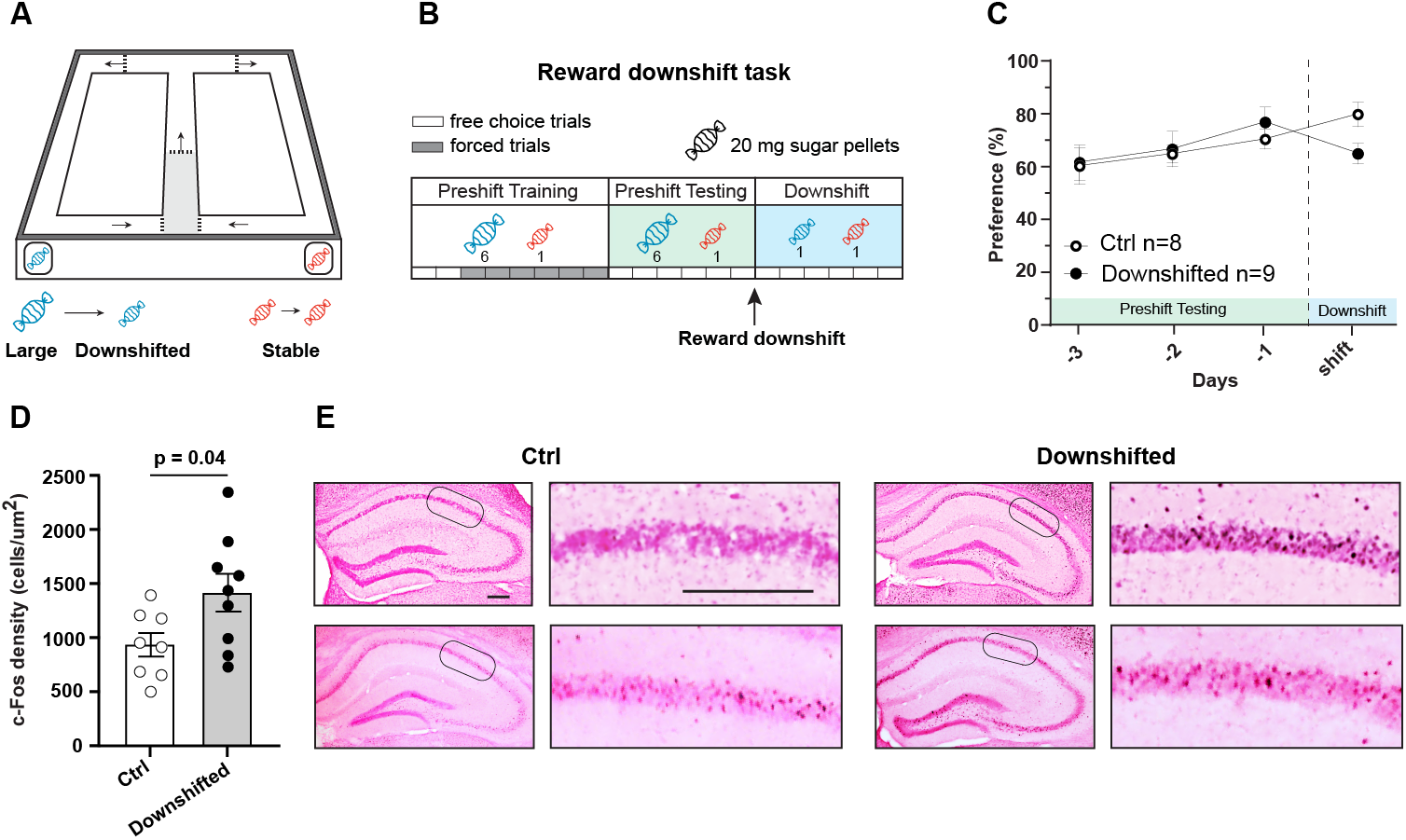
Unexpected reward downshift increases c-Fos recruitment in dorsal CA1. (A) Schematic representation of the figure-8 maze utilized for the instrumental reward downshift task. The maze is equipped with manual doors (dotted lines) and reward locations where sugar pellets are delivered. Mice were held in the delay zone (shaded region) for 15 seconds between trials. (B) The experimental timeline consisted of two main contingencies: Preshift (unequal rewards of 6 mg and 1 mg sugar pellets) and Downshift (the large reward is reduced to 1 mg to match the magnitude of the stable reward). The Preshift phase included a 7-days session training period consisting of both free-choice and guided ‘forced’ trials (grey and white key). During the 3 sessions of Preshift testing and the subsequent Downshift session, only free-choice trials were administered to assess preference. (C) Behavioral summary (n = 8 control mice, n = 9 downshifted mice; mean ± sem). During the Preshift contingency, mice developed a significant preference for the reward location associated with the large magnitude reward, which was subsequently reduced following the unexpected reward downshift. Preference for the large reward was defined as ≥ 70% free-choice trials toward the larger reward port on 2 of 3 consecutive days. (D) Quantification of c-Fos density (coordinates from bregma −1.6 to −1.8 mm AP) in the dorsal CA1 of Control and Downshifted groups. A Mann-Whitney U test revealed a significant increase in c-Fos density in the Downshifted group compared to the Control group (U= 14, p= .036), providing evidence of enhanced network engagement during the violation of reward expectancy. (E) Representative micrographs of dorsal hippocampal sections showing c-Fos immunostaining. The first column displays two examples from the Control group with corresponding high-magnification insets of the CA1 pyramidal layer (magnified areas indicated by rounded rectangles). The second column displays two examples from the Downshifted group with corresponding CA1 magnifications. Scale bars represent 200 μm.

**Extended Data Figure 2:**
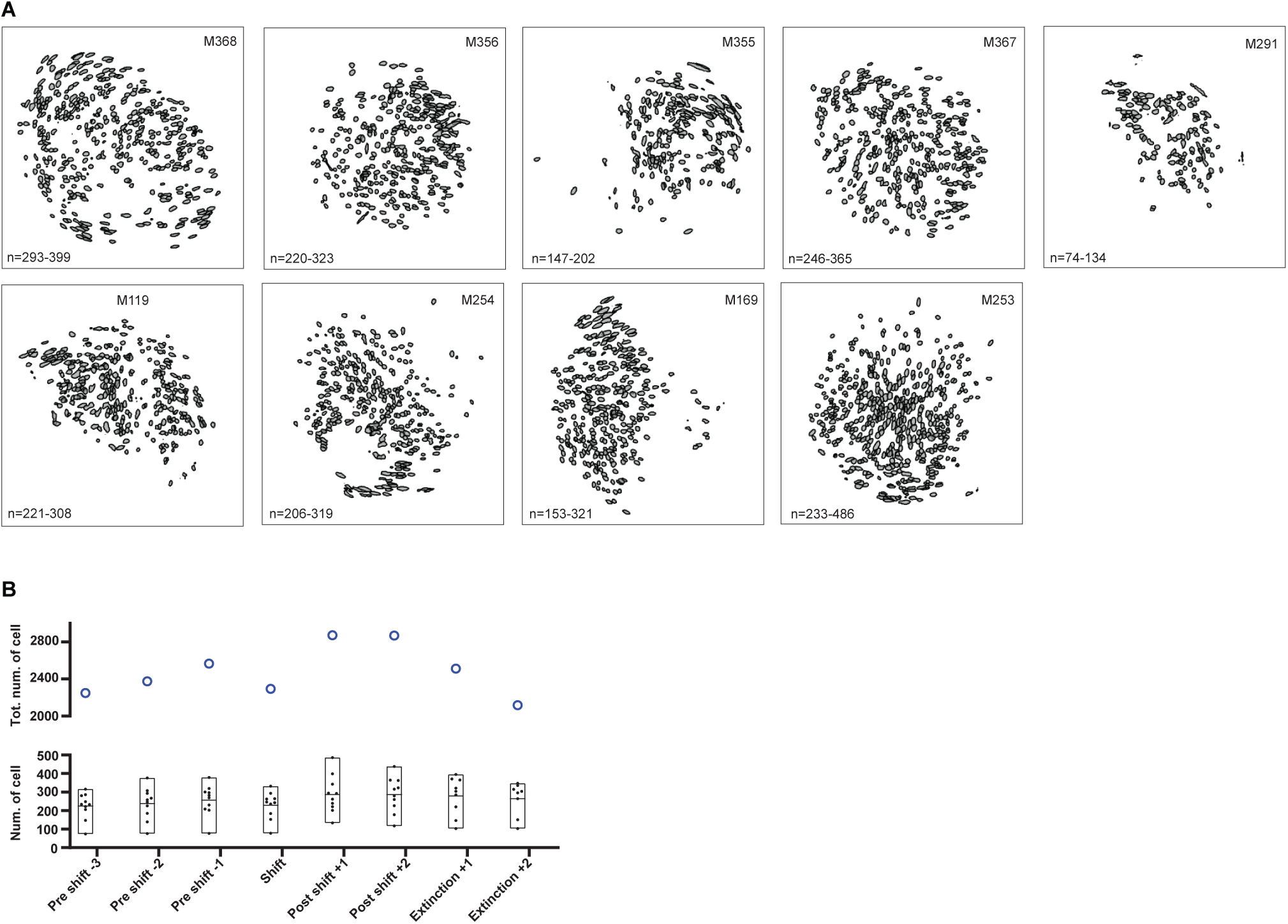
Spatial footprints of extracted neurons from all mice. (A) Spatial footprints of extracted neurons from all mice (n = 9). Numbers indicate the minimum and maximum number of extracted units across sessions. (B) Quantification of the number of neurons extracted from each animal across all imaging sessions (bottom), and the total number of extracted neurons pooled across all mice (top).

**Extended Data Figure 3:**
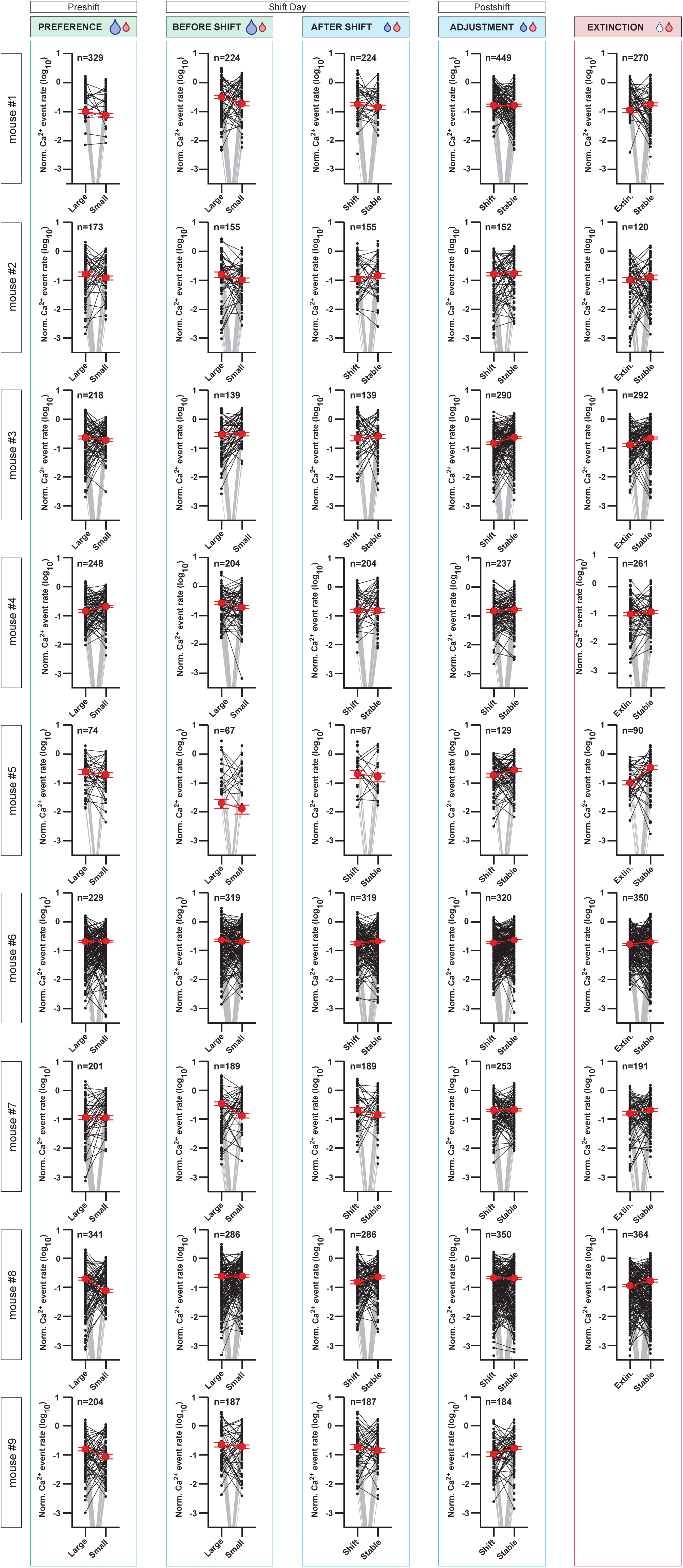
Calcium events rate in individual mice. Normalized calcium event rates at reward locations across task phases. Each panel shows individual neuron activity for a single mouse during the reward downshift task. Red dots and lines indicate the mean normalized event rate across all recorded neurons. Grey lines connect neurons that were active at only one of the two reward locations. Note that such neurons go off scale because of the log scale.

**Extended Data Figure 4:**
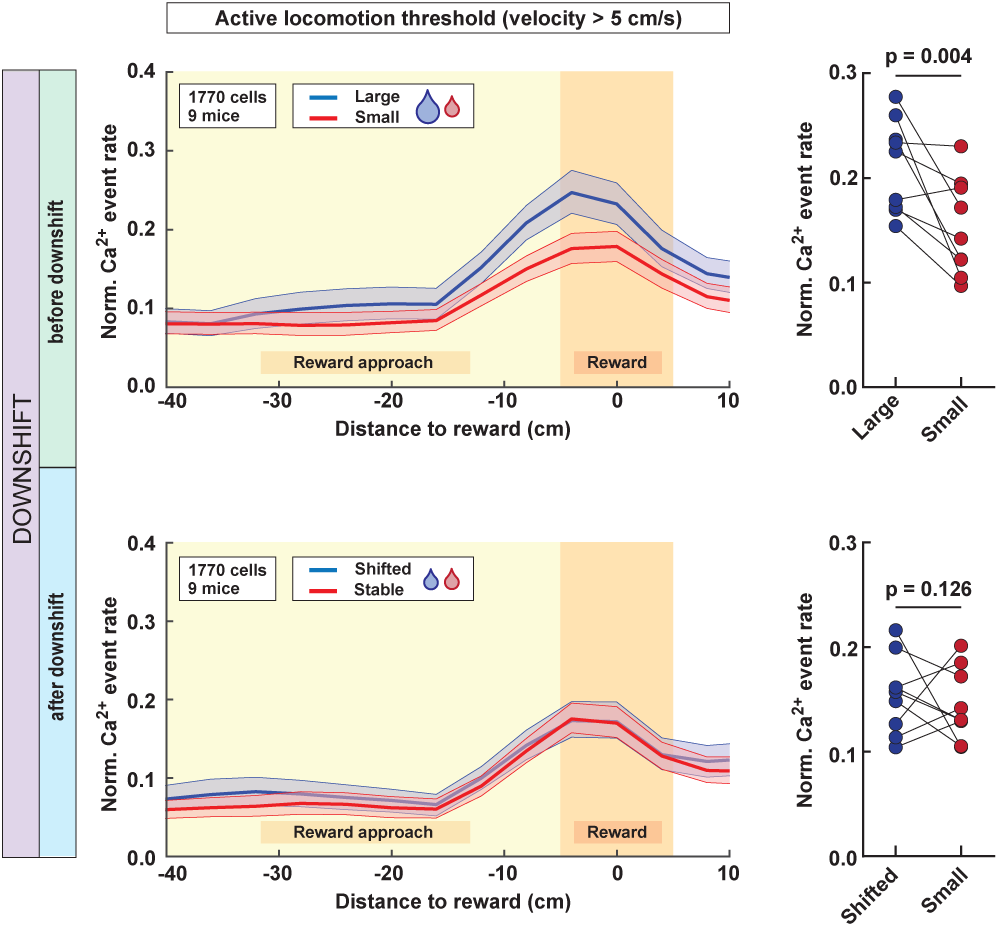
Preservation of reward-magnitude population code at stringent locomotion criteria (>5 cm/s). Trajectory-dependent neuronal activity during Downshift session. (Left) The binned normalized calcium event rate during running for the reward arm (“Reward approach”) and reward zone (“Reward”) during forced choice trials, pooled across all neurons from all mice (n = 9). Blue traces correspond to turns toward the large/shifted reward, and red traces correspond to turns toward the small/stable reward. Shaded areas represent 95% confidence intervals. (Right) Mean normalized calcium event rate at reward locations averaged across neurons for each mouse. P-values shown for generalized linear mixed effects model comparing activity across rewards.

